# Evidence of silent nociceptor activation during the formalin test in rats

**DOI:** 10.64898/2026.11.10.765590

**Authors:** Elise A. Ajay, Anthony E. Pickering, James P. Dunham

## Abstract

**Background:** Abnormal ongoing activity in C-mechano-insensitive (C-Mi) nociceptors is seen in patients with chronic pain. These ‘silent’ nociceptors are often under-reported in preclinical pain models – precisely because they aren’t mechanically sensitive. The formalin test is widely used as a preclinical model of persistent sensitization characterised by two behavioural phases, where phase 2 is considered to reflect central sensitisation and the role of peripheral afferent drive in its maintenance is disputed. Increased activity in C-mechanosensitive (C-M) nociceptors has been reported during both phases, but the behaviour of C-Mi nociceptors has not been assessed.

**Methods:** Multi-contact microneurography was conducted from the saphenous nerve of male and female rats. An activity dependent slowing protocol was used to segregate C-fibres, and responses to mechanical stimuli were assessed. Formalin- (2.5%) or saline-evoked activity was then assessed during the two test phases.

**Results:** 168 A- and C-fibres were identified with latency tracking, and classified using activity dependent slowing into C-M and C-Mi nociceptors. ADS-classified C-Mi nociceptors were active in both phase 1 and phase 2 (33.3% and 31.0%, n=18 and n=16, respectively). We find no evidence that formalin mechanically sensitises C-Mi nociceptors or alters the ADS profiles of C-fibres. There was no sex specific difference in C-fibre classes or the effect of formalin.

**Conclusions:** These results indicate the presence of ongoing activity in a substantial population of C-Mi nociceptors in a commonly used persistent pain assay that may play a role in the sensitisation that is seen in phase 2 of the behavioural response.

**Key Points:**

1. Abnormal ongoing activity in silent (also known as C-mechano-insensitive, C-Mi) nociceptors is seen in patients with chronic pain, but the contribution of these important sensory afferents in pre-clinical nociceptive assays, such as the formalin test, is unclear.
2. Using *in vivo* electrophysiology and applying activity dependent slowing and mechanical sensitivity testing, we identify a C-Mi nociceptor population in the saphenous nerve of anaesthetised rats, and track their activity after exposure to subcutaneous formalin or saline injections.
3. We observe that both C-mechano-sensitive and C-Mi nociceptors are active during phase 1 and phase 2 of the formalin test in rats at similar proportions (∼30%).
4. These results support the theory that ongoing nociceptor activity is important in maintaining central sensitisation, as seen in previous studies with humans, and provides clarity regarding pre-clinical nociceptive assays and their relevance to human disease.

## Introduction

Preclinical pain assays are widely used to investigate pain neurobiology and identify potential targets for chronic pain treatment. Reverse translation by seeking features in preclinical models that mirror abnormalities seen in patients may enhance the utility of these assays. This is relevant for spontaneous nociceptor activity, particularly in ‘silent’ nociceptors, which has been reported in a diverse range of chronic pain conditions (Choi et al., 2024). Silent nociceptors, also referred to as C-mechanoinsensitive (C-Mi) nociceptors or Type 1B C-nociceptors, only become active and mechanically sensitive after localised injury (Gold & Gebhart, 2010; Michaelis et al., 1996). Aberrant activity of C-Mi nociceptors in humans has been implicated as driving neuropathic pain (Beisswanger et al., 2025; Ørstavik et al., 2006; Serra et al., 2012), and is also reported in nociplastic pain conditions (Evdokimov et al., 2019; Ribeiro et al., 2025).

The formalin test is a widely used preclinical model of persistent sensitization (Dubuisson & Dennis, 1977). Formalin is injected subcutaneously into the paw of rodents evoking quantifiable nocifensive behaviours. In rodents, the response is biphasic (Tjølsen et al., 1992): phase 1 (5-10 minutes) is thought to be mediated by direct activation of nociceptive afferents by formalin, and phase 2 (∼30 minutes), separated by brief quiescence, is believed to be driven by central sensitisation possibly maintained by ongoing peripheral nerve activity.

*In vivo* electrophysiological studies in rats have shown strong A- and C-fibre drive during the two phases (Dickenson & Sullivan, 1987a, 1987a; McCall et al., 1996; Puig & Sorkin, 1996), and have reported that ongoing peripheral activity is necessary for eliciting phase 2 behaviours (Coderre et al., 1990; Dallel et al., 1995; Dickenson & Sullivan, 1987a, 1987b). These studies report activity in mechanically sensitive C-fibres in both phases, however the contribution of silent nociceptors remains unknown.

Silent nociceptors are often under-reported due to their lack of response to common mechanical search strategies. However, C-Mi can be identified by their pattern of activity dependent slowing (ADS) which has been documented in humans (Schmelz et al., 2000; Serra et al., 2004), pigs (Obreja et al., 2010) and rats (Hoffmann et al., 2015; Serra et al., 2012). ADS is an intrinsic physiological property of axons, whereby repeated action potential discharge reduces conduction velocity, (i.e., slowing). This phenomenon can be revealed by applying electrical stimulation trains to the receptive field. Critically, silent nociceptors have a high degree of slowing at low frequencies of stimulation.

Here we have applied *in vivo* microneurography using multi-contact electrodes (Sales et al., 2022) in the intact saphenous nerve of the rat together with an activity dependent slowing protocol. We show that both C-mechanosensitive (C-M) and C-Mi nociceptors are active during phase 1, and show ongoing spontaneous activity during the phase 2 of the formalin test.

## Materials and Methods

### Animals and ethics

All experiments and procedures conformed to the UK Animals (Scientific Procedures) Act 1986 and were approved by the University of Bristol Animal Welfare and Ethical Review Body. The experimental design and reporting follow the ARRIVE Guidelines. Female (n = 12) and male (n = 13) adult Han Wistar rats (200-420g) were purchased from Charles River Laboratories and housed under a 12 hour light/dark cycle with *ad libitum* access to food and water.

### Surgery

All procedures were conducted under terminal general anaesthesia. Anaesthesia was induced using 5% isoflurane and oxygen (1 L/min) until the pinch withdrawal was lost. Rats were maintained areflexic on 1.5-3% isoflurane and oxygen (0.5-1 L/min). Core body temperature was maintained at 36°C using a heat mat and rectal thermometer throughout surgery and recordings.

The rat was positioned supine and the medial side of the right hind limb was shaved from ankle to hip. An incision was made on the medial aspect of the hind limb from approximately the inguinal ligament to the tibiofemoral joint. Silk suture was tied around the patellar ligament and passed through the skin to externally rotate and stabilise the hind limb. In preparation for cutaneous electrical stimulation, a steel needle electrode was inserted subcutaneously into the plantar surface of the paw prior to securing the paw in position. The skin was separated from underlying tissues using blunt dissection and sutured onto a metal loop to create a pool that was filled with saline.

The saphenous nerve proximal to the knee was gently separated from the underlying tissues. With fine forceps, the outer layers of the nerve sheath were dissected from the nerve, leaving the last layers of the epineurium intact. The exposed nerve was lifted and a small block of 5-10% agar (∼1 x 2 mm) was positioned underneath. An Ag/AgCl reference electrode (EP1, World Precision Instruments; USA) was positioned near to the recording site and secured. The saline in the bath was removed. The bath was then filled with 3% agar in 0.9% saline to stabilize the nerve and keep the tissues hydrated.

### Solutions for injection

Subcutaneous injections made during the recording protocol were either equivalent injection volumes (30 µl) of formalin 2.5% (333 mM) in saline (VWR, USA) or sterile saline (0.9%). These solutions were prepared prior to all experiments and aliquoted into coded tubes for blinding. A 100 µl Hamilton syringe fitted with a 12 µl volume catheter fitted with a 27 gauge steel needle end was used to deliver the solution during recordings. The catheter was flushed with saline then prefilled with 10 µl of the solution, intentionally leaving 2 µl of saline at the tip to avoid interference in the initial parts of the experiment.

### Nerve recordings

A 32 channel microelectrode array probe (A1x32-Poly3-10mm-25s-177-A32; NeuroNexus, USA) was used to record neural activity from the saphenous nerve. Before experiments, the electrode impedances were tested in saline to confirm functionality. Recordings were acquired via an Intan RHD2232 amplifier chip (USA) and Open Ephys acquisition system (version 0.6.6; USA) using a 30 kHz sampling rate. Using a micromanipulator, the probe was inserted through the solidified 3% agar into the partially de-sheathed nerve over the agar block. If neural activity was identified on the electrodes, a search for a receptive field was conducted by gentle brushing of the paw and ankle. If there was no spontaneous activity and/or no identifiable receptive field, the probe was reinserted into the nerve in a new position and re-tested to locate an accessible receptive field (Fig. 1a).

**Figure 1:**
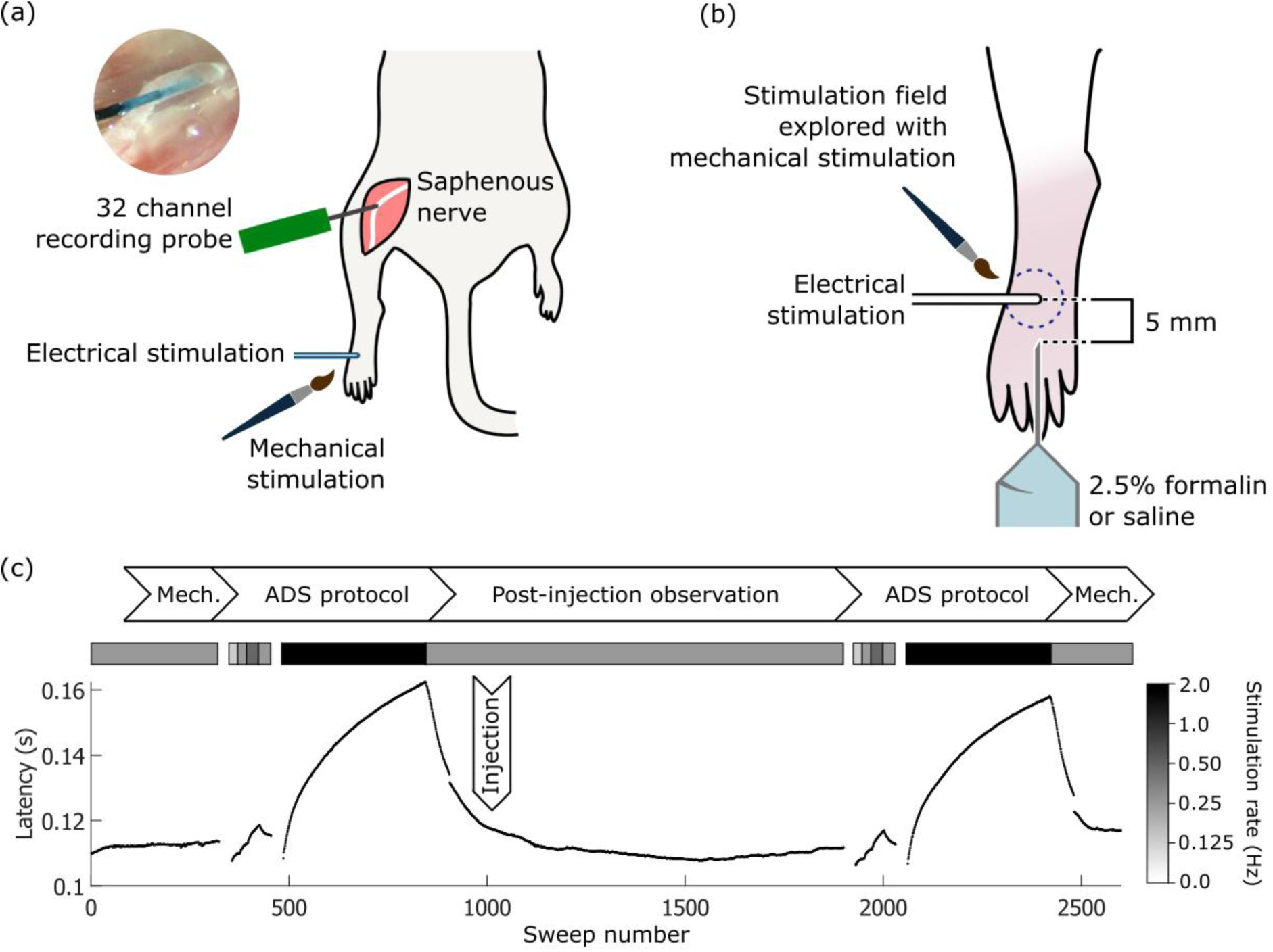
Experimental set up and procedure. (a) A 32 channel recording probe was inserted using a micromanipulator into the saphenous nerve to capture single unit activity on multiple contacts. A receptive field for the electrode was identified on the dorsum of the paw using electrical and mechanical stimulation. (b) A 27 g needle catheter pre-loaded with the treatment (2.5% formalin or saline) was inserted subcutaneously such that the tip of the catheter needle was 5 mm from the stimulation site at the start of the experiment. The stimulation field was explored with mechanical stimulation (brush and von Frey filaments) to identify mechanically sensitive units. (c) An example of a latency track belonging to a C-fibre nociceptor, with the stimulation rate represented above the latency track as coloured bars (legend to the plot right). The protocol ffow chart sits above, and the timing of the injection indicated with an overlaid arrowbox. Each black dot on the plot represents the latency of the electrically-evoked spikes of the identified nociceptor.

### Experimental protocol

Using a steel needle electrode held in a micromanipulator, an electrical search stimulus (3.2 mA, 0.5 ms, 0.25 Hz) delivered from a constant current stimulator (DS3, Digitimer, UK) was used to find putative C-fibre units (based on approximate conduction velocity <1 m/s) in the receptive field using APTrack (Nickerson et al., 2023). These C-fibre units were confirmed by adjusting the electrical stimulus intensity and duration to confirm an all-or-nothing response and latency indicative of C-fibres (Fig. 1b). The position of the needle electrode was carefully marked on the skin for ease of visualisation. This was important to confirm if movement of the electrode relative to the skin occurred during the experiment and to measure the conduction distance at the end of the experiment.

In 19 rats (n = 8 female), the needle of the Hamilton syringe catheter was inserted subcutaneously in the dorsum of the paw such that the tip was approximately 5 mm from the stimulating electrode. A 0.25 Hz electrical stimulus (3.2 mA, 0.5 ms) was applied for 10 minutes to allow the latencies of all units to stabilise. Mechanical sensitivity of units was assessed using the marking technique. The responsiveness of a unit is indicated by a sudden increase in latency to electrical stimulation following mechanical stimulation, followed by a gradual return to the initial latency. Gentle brush and 15 g von Frey were applied to the skin around the stimulating electrode (at least 2 applications of each, as permitted by the anatomy of the rat foot dorsum).

In rats without the Hamilton syringe catheter (n = 6), mechanical testing was similarly assessed with brush and 1-15 g graded von Freys filaments to assess the mechanical thresholds of units.

Mechanical stimulation of the area around the electrical stimulation site typically excited multiple units, the receptive fields for individual units were not mapped.

Following mechanical stimulation, a series of electrical stimuli was applied to characterise the pattern of activity dependent slowing after (Obreja et al., 2010) and as shown in Table 1.

**Table 1:** The activity dependent slowing protocol, showing the series of electrical stimulation frequencies applied to the receptive field to characterise the pattern of slowing of individual nerve fibres.

| **Step** | **Stimulation frequency** | **Number of pulses** | **Duration (seconds)** |
|---|---|---|---|
| 1 | 0 (rest) | N/A | 120 |
| 2 | 0.125 Hz | 20 | 160 |
| 3 | 0.25 Hz | 20 | 80 |
| 4 | 0.5 Hz | 30 | 60 |
| 5 | 0.25 Hz | 30 | 120 |
| 6 | 0 (rest) | N/A | 120 |
| 7 | 2 Hz | 360 | 180 |
| 8 | 0.25 Hz | 60 | 240 |

For animals with the Hamilton syringe catheter inserted subcutaneously, following the initial mechanical and activity dependent slowing characterisation, continuous electrical stimulation was applied at 0.25 Hz. Unit latencies were allowed to stabilise over approximately 5 minutes, before a 30 µl subcutaneous injection of either formalin or saline. Experimenters were blinded to the injectate. Unit latencies were recorded for 60 minutes following the injection, before repeating the activity dependent slowing protocol and then mechanical sensitivity testing (Fig. 1c).

In animals without the Hamilton syringe catheter, a new receptive field was identified and the mechanical testing and activity dependent slowing protocol repeated at the new site. Up to 3 receptive fields per rat were captured.

### Data processing

Recordings were processed offline in MATLAB (R2025a, USA) using a median common average filter and bandpass filter (300-6000 Hz). Electrical artefacts were blanked (i.e., set to the mean) 2 ms before and after electrical stimuli. Putative spikes were identified on each channel as exceeding one standard deviation of the signal RMS, using a peak finding algorithm. Spike latency rasters and waterfall plots for each recording electrode (channel) were generated and used to extract spike latencies of individual units.

The relative latency change during the activity dependent slowing protocols was calculated using automated scripts, except in cases where flip-flop occurred during the protocols, which were manually corrected. Flip-flop, also referred to as hopping, describes an abrupt step in the latency track that occurs when the point of electrical excitation jumps from one point/dendritic branch of the nerve fibre to another (McMahon & Wall, 1987; Torebjörk & Hallin, 1974; Weidner et al., 2003).

Experimenters were blinded to the identity of the injectate during spike and latency track extraction.

### Unit classification

Recordings of A-and C-fibres were separated on the basis of conduction velocity (Fig. 2a). C-fibres were subclassified according to the degree of slowing during the activity dependent slowing protocols (Figs 2b-e), which were then compared to their mechanical sensitivity assessed as described above. Units which slowed initially, then partially recovered during the 2 Hz stimulation were identified as putative sympathetic efferents and incorporated into the C-non-nociceptor class.

**Figure 2.**
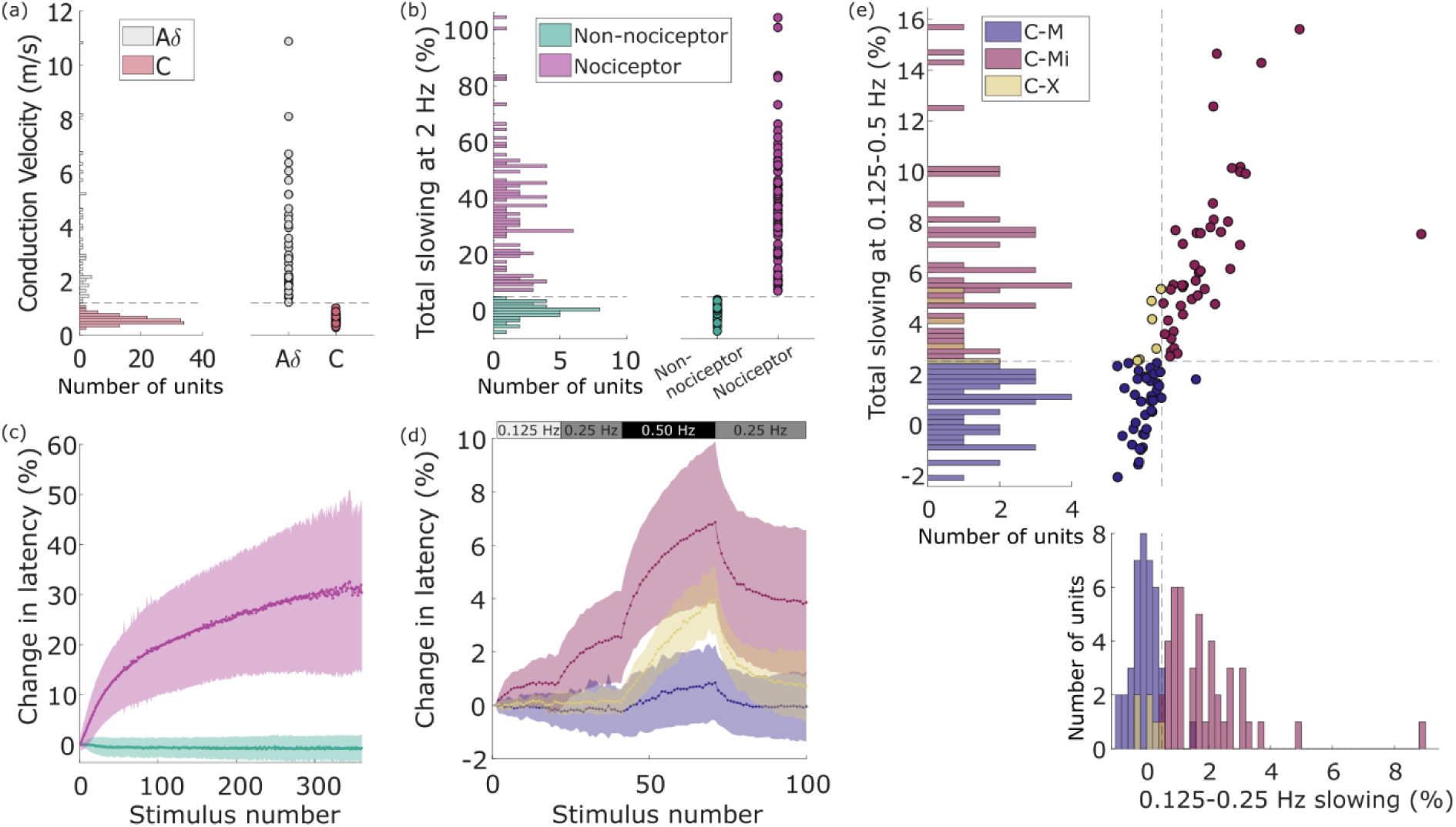
Conduction properties of captured saphenous fibres. (a) Conduction velocities of units split into Aδ (grey) and C-fibre (red) classes using a threshold of 1.2% (grey dotted line). (b) High frequency (2 Hz) activity dependent slowing separates C-fibre nociceptors (magenta) and C-non-nociceptors (green) using a threshold of 5% (grey dotted line). (c) Average high frequency slowing profiles of nociceptor (magenta) and C-non-nociceptor (green) fibres. Lines represent the mean, and shaded region represents the standard deviation. (d) Average activity dependent slowing profiles of the putative mechanically sensitive (C-M; indigo), putative mechanically insensitive (C-Mi; burgundy) nociceptors, and C-X (yellow) nociceptors. Lines represent the mean, and shaded region represents the standard deviation. The stimulation frequencies are presented at the top of the plot. (e) Total slowing during low frequency stimulation protocol, and specifically the slowing during the initial 0.125-0.25 Hz stimulation, reveals two clusters of nociceptors, defining the putative C-M (indigo) and putative C-Mi (burgundy) nociceptors. Units which showed high slowing at low frequencies but less than 0.5% in the 0.125-0.25 Hz phase were labelled as C-X (yellow).

### Formalin test Phase 1 and Phase 2 activity classification

Phase 1 was defined as the 10 minute period immediately following the injection of formalin (or saline), and phase 2 as the period 10-60 minutes after injection. The presence of ongoing activity in a C-fibre resulting from the subcutaneous injection can be inferred by the presence of latency jitter or complete failures to electrically evoke a response because of a refractory period after spike discharge from ongoing activity, which then recovered with time. Ongoing activity during phase 1 was defined as a failure to evoke a spike at the expected latency greater than 8 consecutive sweeps at 0.25 Hz (i.e., 32 seconds) starting at least 12 seconds after the injection to exclude any possible immediate mechanical effect. The duration of phase 1 activity was calculated as the duration of the failure to electrically evoke a response in minutes.

Phase 2 activity was identified by disruptions in the latency tracks, including double spikes, jitter, and/or abrupt or rounded changes in the latency tracks that were not seen on other units, and are indicative of ongoing activity.

Experimenters remained blinded to intervention allocation for the analysis of phase 1 and phase 2 activity.

### Statistics

A Kaplan-Meier survival analysis was applied for mechanical sensitivity comparisons in C-M and C-Mi nociceptors.

Fisher’s exact tests were applied for comparisons of phase 1 and phase 2 activity in formalin- and saline-treated C-nociceptors, and in male and female rats.

A Chi-square test was applied for sex differences in mechanical sensitivity.

All data are reported as means ± standard error mean unless stated otherwise.

## Results

### Afferent Classification

We first used latency tracking and human activity dependent slowing protocols to identify silent C-nociceptors in the rat saphenous nerve. Stable latency recordings were made from 168 well-isolated units in the saphenous nerve using multi-contact electrodes (n = 25 rats, of which n = 12 female). Units were extracted from 31 receptive fields, yielding 5 (3, 7; median and Q1, Q3) units per receptive field. Units were initially segregated by conduction velocity into Aδ fibres (1.2-13.7 m/s) and C-fibres (≤1.2 m/s). This threshold (1.2 m/s) was chosen based on the frequency histogram shown in Fig. 2a which shows a distinct separation between two populations, with the upper limit bounded at 1.2 m/s. This yielded 127 (75.6%) C-fibres and 41 (24.4%) A-fibres (Fig. 2a).

During 2 Hz stimulation of all C-fibres, two populations were apparent, those with minimal increases or decreases in latency, and those that showed a considerable increase in latency by the end of the 2 Hz stimulation. These populations were segregated by the frequency histogram shown in Figure 2B into putative nociceptors (>5% slowing) and putative non-nociceptors (<5% slowing). This identified 89/127 (70.1%) C- nociceptors and 38 (29.9%) as C-non-nociceptors (Figs 2b-c, Table 2).

**Table 2:**
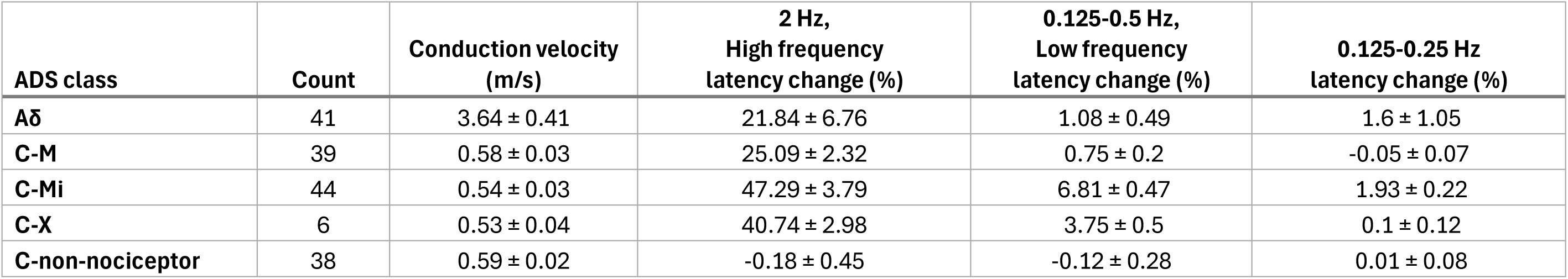
Summary of conduction velocities and latency changes during the activity dependent slowing protocol for saphenous nerve fibre classes.

| **ADS class** | **Count** | **Conduction velocity<br>(m/s)** | **2 Hz,<br>High frequency<br>latency change (%)** | **0.125-0.5 Hz,<br>Low frequency<br>latency change (%)** | **0.125-0.25 Hz<br>latency change (%)** |
|---|---|---|---|---|---|
| **Aδ** | 41 | 3.64 ± 0.41 | 21.84 ± 6.76 | 1.08 ± 0.49 | 1.6 ± 1.05 |
| **C-M** | 39 | 0.58 ± 0.03 | 25.09 ± 2.32 | 0.75 ± 0.2 | -0.05 ± 0.07 |
| **C-Mi** | 44 | 0.54 ± 0.03 | 47.29 ± 3.79 | 6.81 ± 0.47 | 1.93 ± 0.22 |
| **C-X** | 6 | 0.53 ± 0.04 | 40.74 ± 2.98 | 3.75 ± 0.5 | 0.1 ± 0.12 |
| **C-non-nociceptor** | 38 | 0.59 ± 0.02 | -0.18 ± 0.45 | -0.12 ± 0.28 | 0.01 ± 0.08 |

Putative C-nociceptors were further subcategorised by their activity dependent slowing profiles during low frequency (0.125-0.5 Hz) stimulation into two groups: those with a high degree of slowing and those with low slowing, as would be anticipated for C-Mi and C-M populations, respectively (Fig. 2d). To identify this population, total slowing over the entire low frequency stimulation paradigm (0.125-0.5 Hz), was plotted against slowing during the 0.125-0.25 Hz phase. As can be seen in Figure 2E, there are two partially overlapping populations which appear to be segregable by a total low frequency slowing of 2.5%, and by the 0.125-0.25 Hz phase slowing of 0.5%. This revealed 44/89 (49.4%) with a high degree of slowing during low frequency stimulation labelled as putative C-Mi, and 39/89 (43.8%) without identified as putative C-M nociceptors. A small number of C- nociceptors (n = 6/89; 6.7%) sat in an intermediate boundary zone showing a high degree of slowing only during the 0.5 Hz stimulation, and these units were labelled as C-X (Table 2).

Following activity dependent slowing characterisation, we applied mechanical sensitivity testing to validate our assessment. The mechanical sensitivity of units was assessed using the marking technique (see methods) whilst applying gentle brush or von Frey filaments around the electrical stimulation site. In most cases, gentle brush or 15 g von Frey were applied, however in a some cases, the mechanical thresholds were determined by additionally applying 1-8 g von Frey filaments. Of the 168 units that were classified using activity dependent slowing, 11 (6.5%) could not be reliably tracked during the mechanical stimulation period (due to overlapping activity from other units, for example), and were therefore labelled as “unknown”.

Table 3 shows the responses to mechanical stimulation of all units in the putative C-M and C-Mi classes. It further shows the mechanical threshold of those units that were assessed in more detail with graded von Frey filaments. A Kaplan-Meier survival analysis (Fig. 3c), which accounts for the censured data (i.e., the non-responders) demonstrates that when the threshold of units are determined with graded von Frey filaments, putative C-Mi fibres respond to mechanical stimuli less frequently (n=8/15, 53.3%) than C-M fibres (n=9/10, 90.0%). When they do respond, the putative C-Mi fibres have higher thresholds with the median activation threshold for the C-M population being 4 g, compared to 15 g for the C-Mi population. Plotting the proportion of C-nociceptors that responded to the 15 g von Frey filament against their total slowing during the low frequency ADS protocol does not reveal two distinct populations of mechanically sensitive and mechanically insensitive nociceptors. However, the ADS criteria described above are enriched for C-Mi nociceptors in that units that slowed more than 2.5% during low frequency stimulation are more likely to be mechanically insensitive or are activated by more intense mechanical stimulation (Fig. 3d).

**Figure 3.**
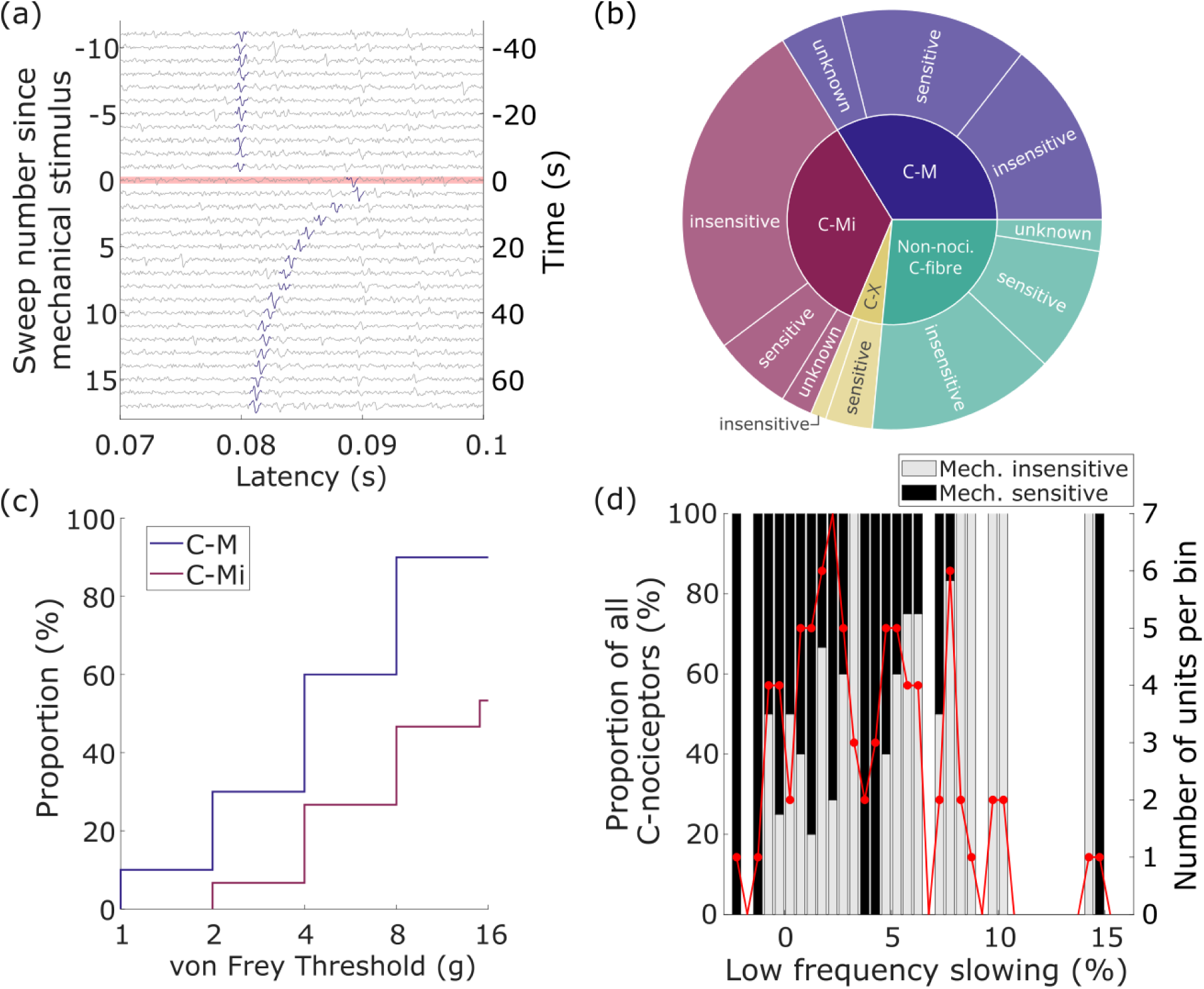
Mechanical sensitivity of identified saphenous nerve C-fibres to brush and von Frey filaments. (a) Example response to 15 g von Frey stimulation of a C-M unit using the marking technique. Red bar indicates timing of application of 15 g von Frey to receptive field. (b) Mechanical sensitivity of all C-fibres. (c) Kaplan-Meier plot showing proportions of units responding to graded von Frey filaments. (d) Proportion of putative C-M and C-Mi fibres responding to brush or von Frey filaments against total slowing during the low frequency activity dependent slowing paradigm. The number of units per histogram bin is plotted on the right axis (red).

**Table 3:**
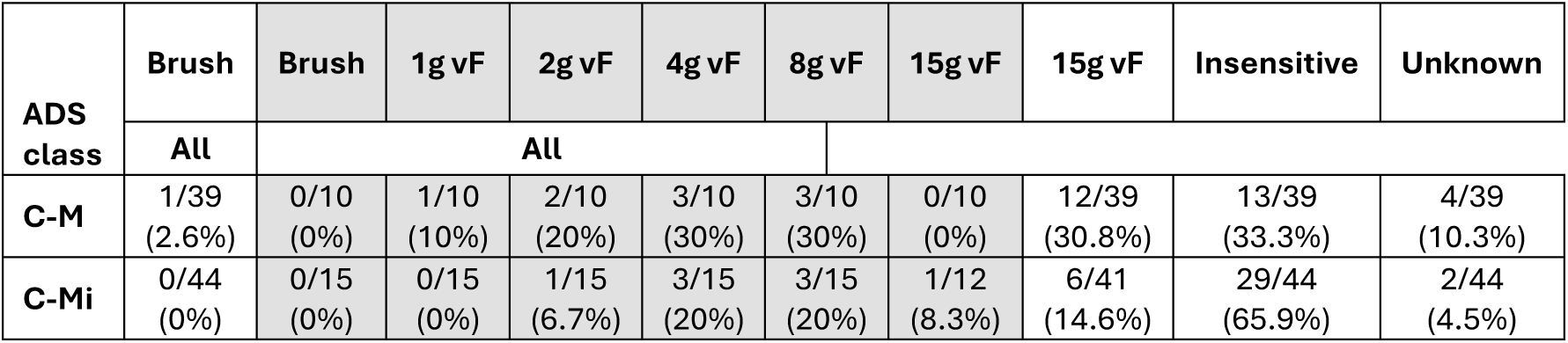
Summary of mechanical sensitivities of fibre classes to brush or von Frey filaments for C-M and C-Mi nociceptors.

At one receptive field, the 15 g von Frey was not applied due to concerns it would displace the receptive field-stimulating electrode position interface. There were 3 C-Mi nociceptors in this receptive field that didn’t respond to the 8 g von Frey filament, and the denominators in Table 3 reflect this.

Due to the small numbers, C-X units were not included in further analyses. The C-fibre populations will be hereon referred to by their activity dependent slowing profile categorisation, with functional (mechanical) characterisation information included where appropriate.

### Formalin exposure – phase 1

Of the 121 characterised C-fibres, 80 received either formalin or saline via subcutaneous injection adjacent to the electrical receptive field (Table 4). The spike latency tracks of each fibre were recorded over the following hour to capture activity during phase 1 (0-10 minutes after injection) and phase 2 (10-60 minutes).

**Table 4:** Number of rats and C-fibres exposed to formalin or saline.

|  | **Rats** |  | **ADS characterised units** |  |  |  |
|---|---|---|---|---|---|---|
| **Treatment** | **Female** | **Male** | **C-Mi** | **C-M** | **C-non-noci.** | **Total** |
| **Formalin** | 5 | 6 | 20 | 20 | 12 | 52 |
| **Saline** | 3 | 5 | 9 | 9 | 10 | 28 |
| **Total** | 8 | 11 | 29 | 29 | 22 | 80 |

Occasionally, abrupt deflections in latency that rapidly recovered (similar to that seen using the marking technique with mechanical stimuli) were observed during the injection, and these were deemed likely to be mechanically-evoked responses and not included as a response in the phase 1 period.

Over the following minutes after injection disruptions in the latency tracks were observed in 31.3% (25 of 80) of C-fibres. The remainder showed no change in their latency tracks supporting the principle that stable intraneural recordings were maintained. Phase 1 activity in isolated units was typically observed as an increase in latency beginning minutes after injection followed by a complete break in the unit latency tracks reflecting a run of failures of electrically evoked spikes, and then an eventual return of firing to baseline latency (Fig. 4a). This behaviour was more common in formalin-treated C-fibres (14/47; 29.8%) compared to saline-treated C-fibres (2/27; 7.4%) (p < 0.05; Fisher’s exact test). Specifically, formalin-evoked responses were observed in 8/20 C-M and 6/18 C-Mi nociceptors (Table 5). Of these 6 formalin-activated ADS classified C-Mi nociceptors, 5 were mechanically insensitive to mechanical sensitivity testing, and only one responding to 15 g von Frey filament. These findings confirm that formalin activates C-M nociceptors and C-Mi nociceptors to a similar degree.

**Figure 4.**
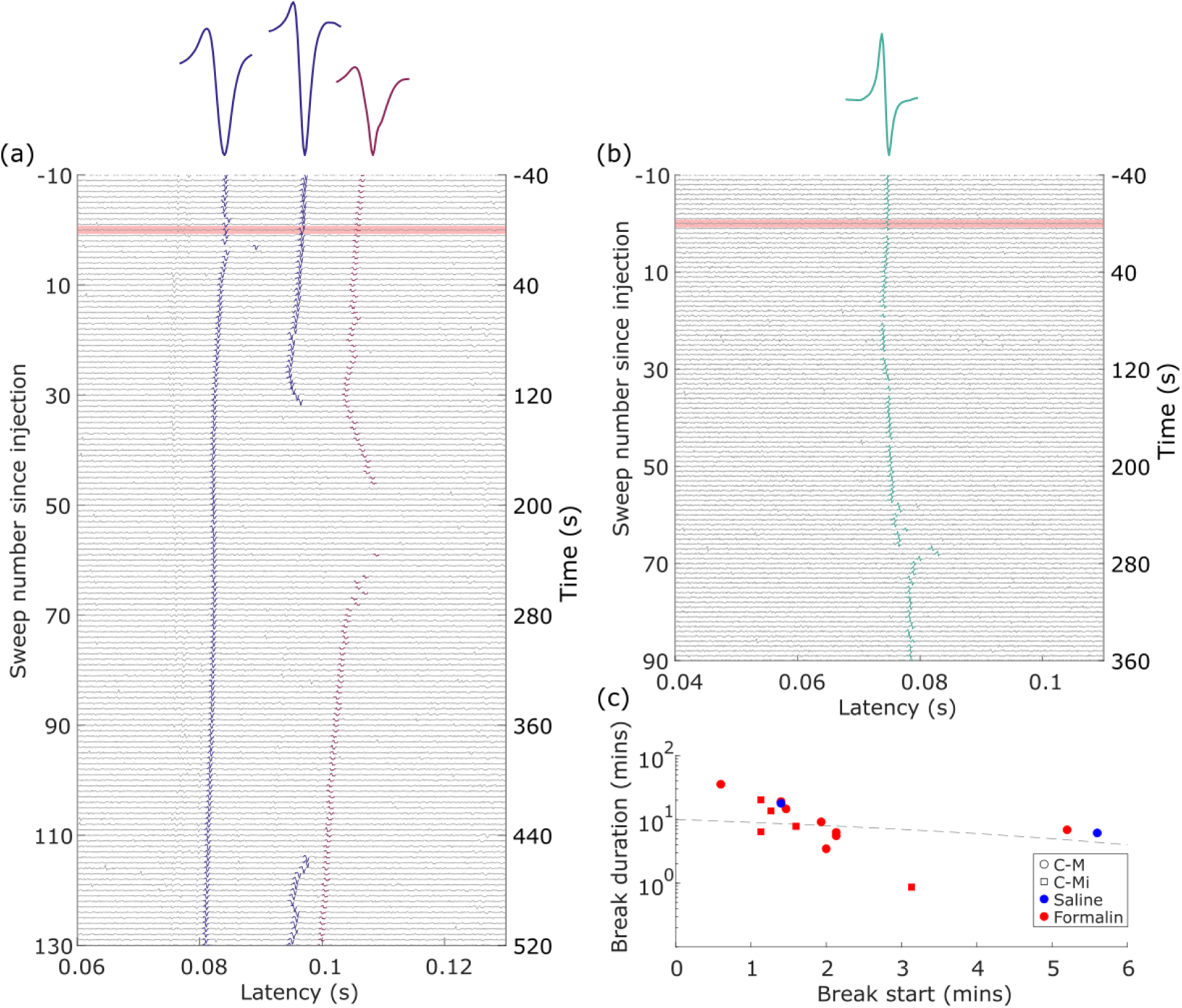
Subcutaneous injection of formalin is associated with disruptions in the latency tracks during the phase 1 period. (a) Example activity in phase 1 period of 3 C-fibres including 2 C-M fibres (blue) and 1 C-Mi fibre (burgundy). Robust increases in activity during phase 1 appears as a break in the latency tracks for the two longest latency (right most) units, and is observed minutes after subcutaneous injection of formalin (red bar) adjacent to the receptive field. (b) Example activity in phase 1 period of a single C-non-nociceptor C-fibre. In this case, activity presents as a deffection in the latency tracks. (c) Time and duration of the break in the latency tracks for formalin-(red) and saline-treated (blue) units. The grey dotted line shows the transition from the phase 1 period to the phase 2 period (i.e., 10 minutes), thus the activity of units above the line extended into the phase 2 period. Symbols indicate unit class (see legend).

**Table 5:** Number of units in each ADS class which showed signs of activity during phase 1 (P1) or phase 2 (P2) in response to formalin or saline injections. Population numbers are reduced during P2 due to ‘loss’ of units during P1, i.e., units stopped responding to electrical stimulation.

|  | *Formalin-exposed units* |  |  | *Saline-exposed units* |  |  |
|  | **P1** |  | **P2** | **P1** |  | **P2** |
| **ADS class** | **Break/Total (%)** | **Deflection/Total (%)** | **Active/Total (%)** | **Break/Total (%)** | **Deflection/Total (%)** | **Active/Total (%)** |
|---|---|---|---|---|---|---|
| **C-M** | 8/20 (40%) | 0/20 | 6/19 (31.6%) | 2/9 (22.2%) | 1/9 (11.1%) | 0/9 |
| **C-Mi** | 6/18 (33.3%) | 2/18 (11.1%) | 5/16 (31.3%) | 0/9 | 3/9 (33.3%) | 0/9 |
| **C-non-nociceptor** | 0/9 | 2/9 (22.2%) | 1/7 (14.3%) | 0/9 | 0/9 | 1/9 (11.1%) |
| **Total** | 14/47 (29.8%) | 4/47 (8.5%) | 12/42 (28.6%) | 2/27 (7.4%) | 4/27 (14.8%) | 1/27 (3.7%) |

An abrupt or rounded deflection in the latency tracks, but no break, was observed after injection in other C-fibres for both formalin (n = 4; 2 C-M, 2 C-non-nociceptors) and saline (n = 4; 3 C-Mi, 1 C-M) (Fig. 4B; Table 5), presumably reflecting a delayed mechanical effect.

Four formalin-treated C-fibre units stopped responding to electrical stimulation during the phase 1 period and were ‘lost’, including 1 C-Mi nociceptor, 2 C-M nociceptors, and 2 C-non-nociceptors (Table 5).

Considering only the unit tracks that were broken, the onset of phase 1 activity (i.e., the beginning of the break in the latency tracks) following formalin or saline injection was 2.0 ± 0.1 minutes (n = 14) and 3.5 ± 1.5 minutes (n = 2), respectively (Fig. 4c). The duration of the phase 1 activity (i.e., length of the break in the latency tracks) was 10.8 ± 0.7 minutes and 12 ± 4.1 minutes for formalin- and saline-treated units, respectively. For many of these units, activity that began in phase 1 extended into phase 2 (n = 9/16, of which n = 7 formalin; Fig. 4c). These units are counted as having phase 1 activity and are not counted again in the phase 2 results below. Figure 4c indicates that there is an inverse relationship between the delay to the activity, and the duration of the break, an observation that could be consistent with an effect of proximity of the fibre’s terminals to the injection location, i.e., the closer the fibre terminals to the injection site, the quicker the onset and longer duration of activation of the fibre.

### Formalin exposure – phase 2

The spike latency tracks in the 10-60 minutes following subcutaneous injection were screened for the presence of abnormal behaviour indicative of spontaneous bursts or double spikes (e.g., abrupt or rounded deflections in latency) (see Methods) (Fig. 5). Formalin was significantly more likely to produce these abnormal behaviours in C-fibres after injection compared to saline (28.6% and 3.6%, respectively; p < 0.05, Fisher’s exact test) (Table 5).

**Figure 5.**
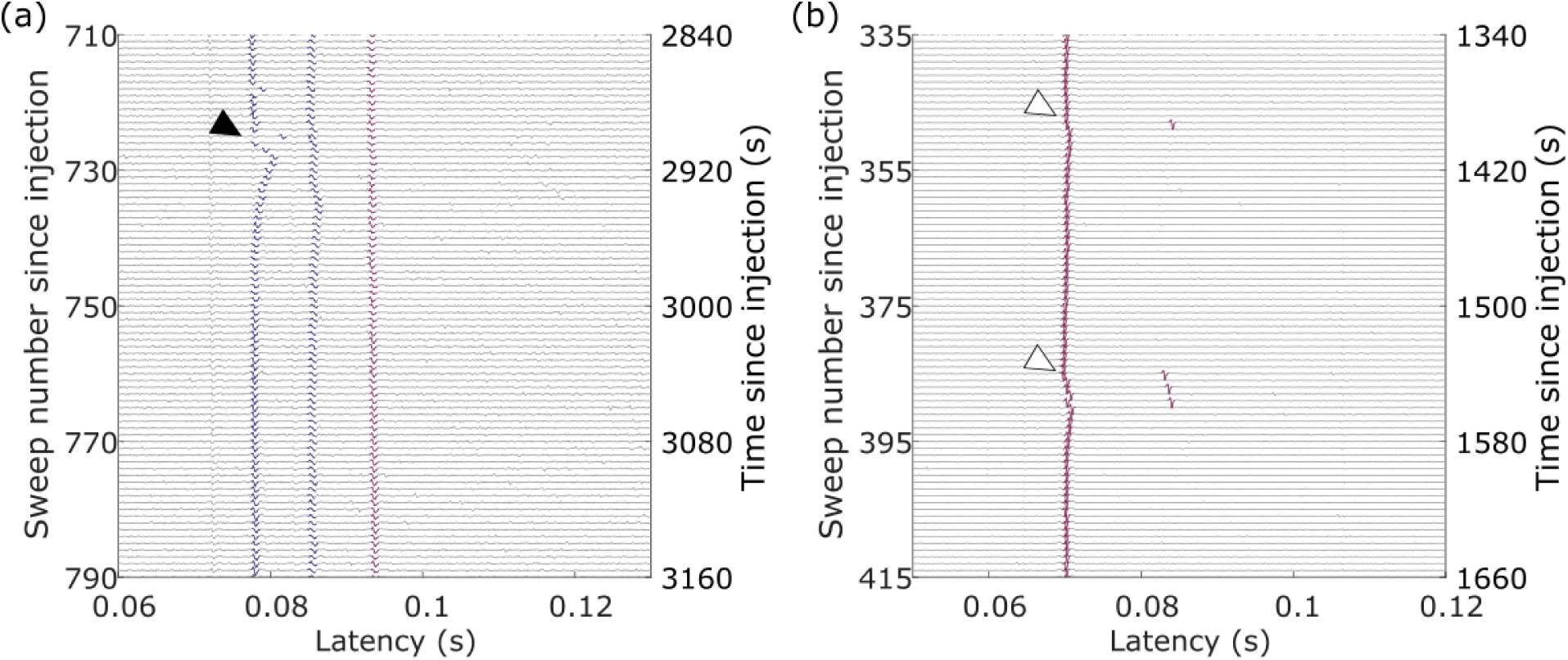
Phase 2 activity in units is observed as abnormalities in the latency tracks. (a) Example of rounded deffections (black triangle) in latency tracks indicative of spontaneous activity. This example shows the same three units from Fig 4a. (b) Example of double spikes (white triangles) causing disruptions, including concomitant increases in latency consistent with activity dependent slowing following double firing, in the latency tracks of an ADS-classified C-Mi nociceptor.

Four units that were active in phase 1 following formalin injection, but then became quiescent, were active again in phase 2, including 3 C-M nociceptors and 1 C-Mi nociceptor. The onset of phase 2 activity was delayed by 20.2 ± 6.5 minutes after the approximate end of phase 1 activity (i.e., return of latency tracks after breaking), indicating an interphase period. None of the saline-treated nociceptor C-fibres were active in both phases.

An additional eight units that weren’t active in phase 1 showed indications of ongoing activity in phase 2 after formalin exposure (Table 5). These units were comprised of 3 C-M nociceptors, 4 C-Mi nociceptors, and 1 C-non-nociceptor. In comparison, in the saline group, no nociceptors and only 1 C-non-nociceptive showed indications of ongoing activity.

In total, 6 (31.6%) C-M and 5 (31.3%) C-Mi nociceptors showed abnormalities in their latency tracks in phase 2 after exposure to formalin. This is in contrast to saline-treated C-M and C-Mi nociceptors, which showed no abnormalities. These five formalin-treated C-Mi nociceptors had activity dependent slowing to low frequency stimulation between 5 and 16%, and did not respond to mechanical stimulation, excepting one whose mechanical sensitivity was “unknown”.

### Formalin effects on activity dependent slowing profiles

Approximately 60 minutes following the subcutaneous injection, activity dependent slowing protocols were repeated. Of the initial 80 C-fibres treated with formalin or saline, the activity dependent slowing profiles of 62 units could be reassessed after the 1 hour observation period (Fig. 6).

**Figure 6:**
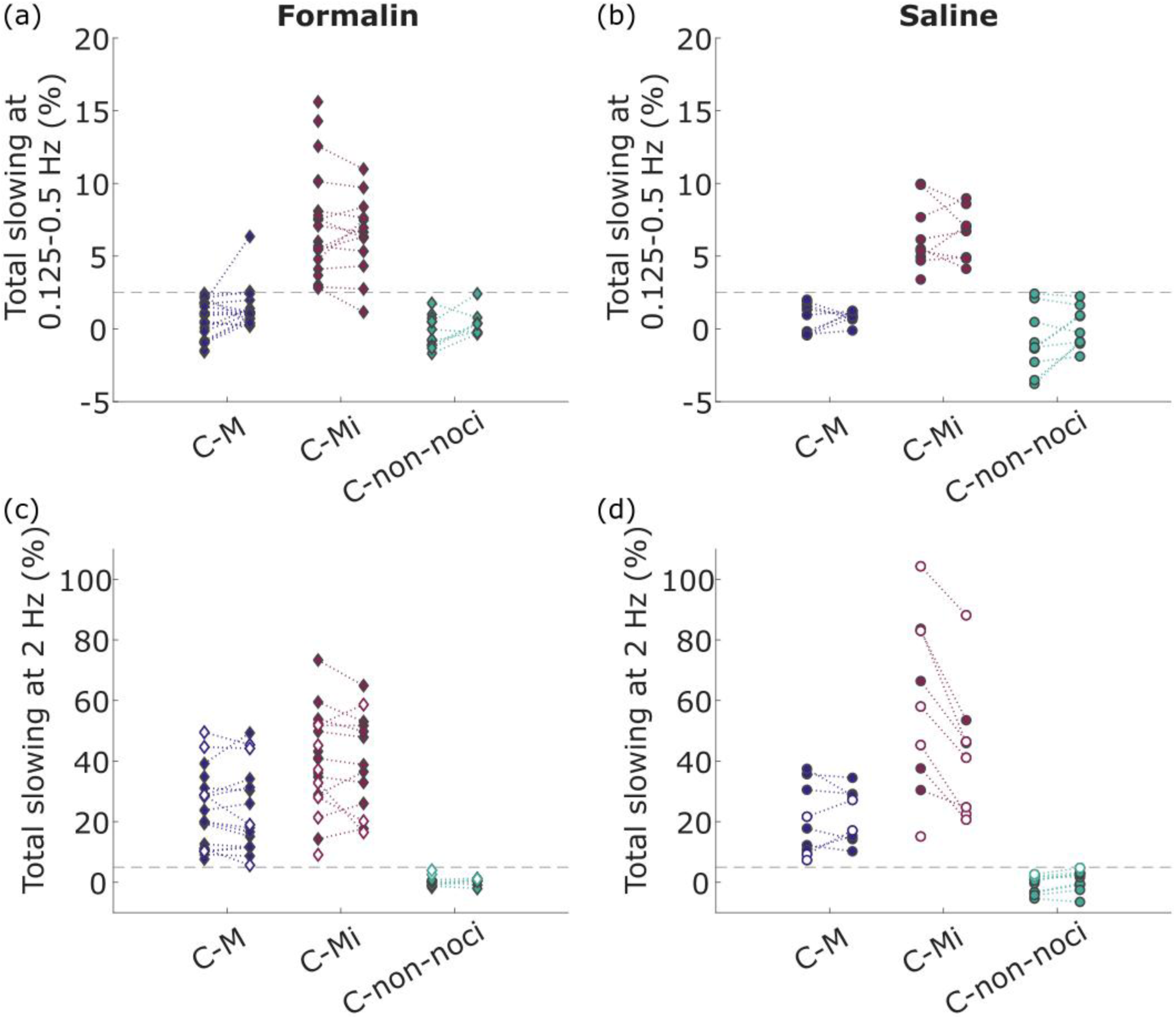
Changes to activity dependent slowing following injection with formalin (left) or saline (right).(a) Change in low frequency slowing (0.125-0.5 Hz) after formalin injection. (b) Change in low frequency slowing after saline injection. (c) Change in high frequency slowing (2 Hz) after formalin injection. (d) Change in high frequency slowing after saline injection. (a) & (b) Dotted line indicates 2.5%. (c) & (d) Dotted line indicates 5%. Open symbols indicate that the unit did not respond to each electrical stimulation (blocking/failure), which will reduce the total activity dependent slowing reached for that unit.

The activity dependent slowing profiles of units were stable when comparing before and after injection, and 60/62 (95.5%) of C-fibres did not change classification. This suggests that activity dependent slowing is a reliable characteristic of C-fibres even after formalin injection. The units which changed classification (n = 2) were both formalin-treated C-nociceptors, and included a C-M and a C-Mi. For these two units, their high frequency (2 Hz) latency change still indicated that the units were nociceptors (>5% increase in latency), however, the low frequency (0.125-0.5 Hz) latency change was altered such that their nociceptor subtype classification also changed. The C-M became a C-Mi nociceptor, and the C-Mi became a C-X.

Interestingly, C-fibre latency tracks for four units appeared only after injection; three after formalin, and one after saline. These units’ activity dependent slowing profiles at the end of the experiment revealed that 2 of the 3 that appeared after formalin were C-Mi nociceptors, and the other a C-non-nociceptor. When these C-Mi fibres were tested at the end of the experiment, they did not appear to be mechanically sensitive. The single unit that appeared after saline injection was a C-non-nociceptor.

### Formalin effects on mechanical sensitivity

Of the 34 C-fibres that were mechanically assessed before and after formalin injection, 10 (29.4%) changed mechanical sensitivity (Table 6). Of these, 5 were sensitised (i.e., previously insensitive now responding to von Frey filaments or brush, or initially only responding to 15 g von Frey filament but now also responding to brush), and 5 lost their previous sensitivity or increased their threshold to mechanical stimuli. In comparison, in the saline injected group only 3/27 (11.1%) units changed their mechanical sensitivity: 2 were sensitised, and 1 was desensitised. Importantly, only 1 of 11 C-Mi nociceptors became sensitised after formalin injection, lending little evidence that formalin exposure mechanically sensitises C-Mi nociceptors in this experimental protocol.

**Table 6:** Change in mechanical sensitivity following formalin injection according to ADS class.

| **ADS class** | **No change** | **Desensitised** | **Sensitised** |
|---|---|---|---|
| **C-M** | 9/14 (64.3%) | 3/14 (21.4%) | 2/14 (14.3%) |
| **C-Mi** | 8/11 (72.7%) | 2/11 (18.2%) | 1/11 (9.1%) |
| **C-non-nociceptor** | 4/6 (66.7%) | 0/6 | 2/6 (33.3%) |
| **Total** | 21/31 (67.7%) | 5/31 (16.1%) | 5/31 (16.1%) |

### Differences between sexes

There was no significant difference in the proportions of C-M, C-Mi, and C-non-nociceptor fibres between males and females (p = 0.18, chi-square test; Table 7). There was also no significant difference in the prevalence of mechanical sensitivity within these classes (p > 0.18 for each class; Table 7), nor in total (p = 0.6946; chi-square test; Table 7).

**Table 7:**
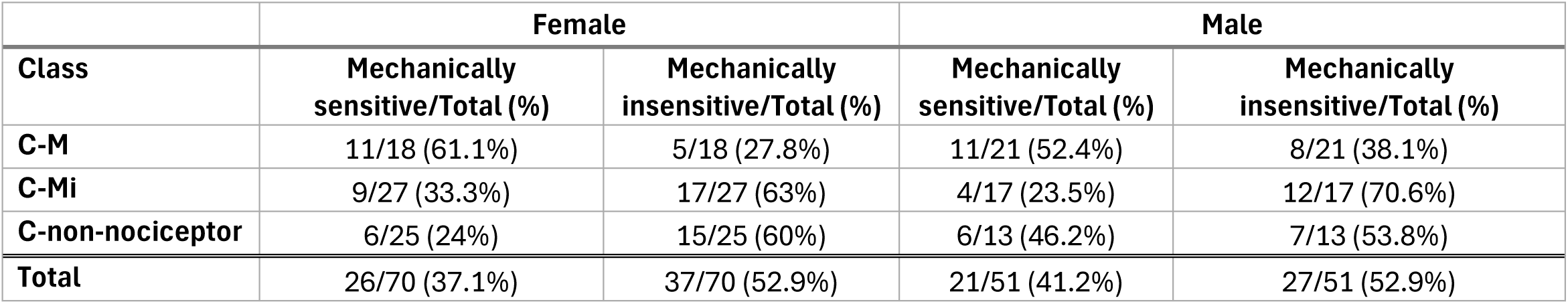
Mechanical sensitivity or insensitivity of saphenous nerve C-fibres in female and male rats according to ADS class.

Latency track disruptions indicating activity following formalin (or saline) injection were equally likely to occur in both sexes (Table 8) (p > 0.05 for P1 breaks and P2 active C-fibres for formalin and saline; Fisher’s exact test).

**Table 8:** Proportion of C-fibres activated by formalin or saline in female and male rats during phase 1 (P1) or phase 2 (P2). C-fibres are grouped according to ADS class.

| **Formalin** |  |  |  |  |  |  |
|---|---|---|---|---|---|---|
|  | **Female** |  |  | **Male** |  |  |
|  | *P1* |  | *P2* | *P1* |  | *P2* |
| **Class** | Break/Total (%) | Deflection/Total (%) | Active/Total (%) | Break/Total (%) | Deflection/Total (%) | Active/Total (%) |
| **C-M** | 3/6 (50%) | 0/6 | 5/6 (83.3%) | 5/14 (35.7%) | 0/14 | 1/13 (7.7%) |
| **C-Mi** | 2/8 (25%) | 1/8 (12.5%) | 2/6 (33.3%) | 4/10 (40%) | 1/10 (10%) | 3/10 (30%) |
| **C-non-nociceptor** | 0/3 | 1/3 (33.3%) | 0/2 | 0/6 | 1/6 (16.7%) | 1/5 (20%) |
| **Total** | 5/17 (29.4%) | 2/17 (11.8%) | 7/14 (50%) | 9/30 (30%) | 2/30 (6.7%) | 5/28 (17.9%) |
| **Saline** |  |  |  |  |  |  |
|  | **Female** |  |  | **Male** |  |  |
|  | *P1* |  | *P2* | *P1* |  | *P2* |
| **Class** | Break/Total (%) | Deflection/Total (%) | Active/Total (%) | Break/Total (%) | Deflection/Total (%) | Active/Total (%) |
| **C-M** | 0/3 | 0/3 | 0/3 | 2/6 (33.3%) | 1/6 (16.7%) | 0/6 |
| **C-Mi** | 0/6 | 2/6 (33.3%) | 0/6 | 0/3 | 1/3 (33.3%) | 0/3 |
| **C-non-nociceptor** | 0/4 | 0/4 | 1/4 (25%) | 0/5 | 0/5 | 0/5 |
| **Total** | 0/13 | 2/13 (15.4%) | 1/13 (7.7%) | 2/14 (14.3%) | 2/14 (14.3%) | 0/14 |

## Discussion

In this study we demonstrate that low and high frequency activity dependent slowing protocols, used in human studies, can be applied to rat sensory afferents to differentiate C-nociceptor classes, identifying a population that is similar to human silent nociceptors. We then applied this observation to identify sensory afferent activity in the two phases of the formalin behavioural test. This demonstrated that around 30% of both C-Mi nociceptors and C-M nociceptors show evidence of formalin-induced spontaneous activity in both phases.

### Identifying functional classes of C-fibres in the saphenous nerve population

To differentiate the populations of sensory afferents in the rat saphenous nerve, we applied activity dependent slowing protocols and mechanical sensitivity testing. During 2 Hz continuous stimulation, a subset of C-fibres exhibited progressive activity dependent slowing, consistent with previous observations that C-nociceptors exhibit substantially greater activity dependent slowing than C-non-nociceptors in rats, pigs, and humans (Obreja et al. 2010, Serra et al. 1999). Prior studies in rats have used >10% slowing during 3 mins of 2 Hz stimulation as the defining criterion to distinguish nociceptive and C-non-nociceptors, based on human experiments (Hoffmann et al., 2015; Serra et al., 2012). After exclusion of putative C-sympathetic efferent units, which slow and then recover during 2 Hz stimulation (Campero et al., 2004), we found a bimodal separation of responses with a cut off at 5%. This value was taken forward based on the empiric data as opposed to utilising cut offs generated in other species. Using this ADS criterion, the majority (70%) of the units we recorded were C-nociceptors, and this is consistent with the literature for foot innervating sensory afferents (Leem, Willis, & Chung, 1993; Leem, Willis, Weller, et al., 1993; Lynn & Carpenter, 1982).

Repetitive low frequency stimulation has also been used to segregate C-nociceptors into functional classes according to their activity dependent slowing. In humans and pigs, greater slowing during low frequency stimulation is strongly associated with a mechanically insensitive phenotype, or, where tested, very high mechanical thresholds (Obreja et al., 2010; Serra et al., 1999; Werland et al., 2021). Our findings support this relationship but demonstrate less complete separation than previously reported in human, pig and rat (*in vitro*) preparations (Hoffmann et al., 2015; Obreja et al., 2010; Serra et al., 1999). Using activity dependent slowing to classify fibres, 29/44 (65.9%) of C-Mi nociceptors were mechanically insensitive, while those that responded to mechanical stimulation exhibited substantially higher mechanical thresholds than fibres classified as C-M. Potential explanations for this observation include, firstly that the segregation between these populations in the rat is not as complete as in the pig and humans.

Secondly, the ADS-classified C-Mi population may also include very-high-threshold (VHT) C-nociceptors, that are sparse in humans, but have indistinguishable ADS profiles to C-Mi nociceptors and mechanical thresholds of ≥150 mN (roughly equivalent to 15 g von Frey) (Werland et al., 2021). Additionally, it is possible that C-Mi afferents, with their larger receptive fields, were sensitised by recurrent stimulation, which is a recognised phenomena in this fibre population in people (Schmidt et al., 2000).

Interestingly, the total high frequency slowing thresholds for separation of C-nociceptors and C-non-nociceptors, and the low frequency slowing threshold for separation of C-Mi and C-M nociceptors were half the thresholds for humans and pigs. The thresholds presented in this study are based on the histogram peaks of the population, although there is no clear dichotomous split in the ADS profiles between the C-M and C-Mi nociceptors and that the cut off enriches for each population but in our hands does not provide an absolute discrimination. This spread of properties of nociceptors echoes the recently reported tiling of mechanical (and thermal) response profiles across genetically identified subtypes in mice (Qi et al., 2024). Further studies using genetic targeting techniques such as optogenetics will be valuable for validating these findings.

The apparent discrepancy between ADS and functional classifications may also be partially explained due to methodological differences. The multi-contact approach enables multiple afferents to be assessed simultaneously. Exploiting this advantage necessitates balancing detailed receptive field and mechanical sensitivity mapping, against the potential for recurrent and potentially noxious stimulation applied to overlapping receptive fields. Thus, not every C-M unit had a response to mechanical stimulation, likely reflecting an absence of stimulation of the most sensitive regions of the receptive field, which are known to be small (Andrew & Greenspan, 1999; Boada et al., 2012). Indeed, when using the 15 g von Frey alone to survey mechanical sensitivity, only 33% of ADS classified C-M fibres were found to respond, whereas 90% were found to respond during graded mechanical thresholding.

### Formalin phases

Our results show that phase 1 of the formalin test robustly activated C-nociceptors, consistent with previous literature (McCall et al., 1996; Puig & Sorkin, 1996). These previous studies have not, however, differentiated between activity evoked by formalin in the mechanically insensitive vs mechanically sensitive C-nociceptors. Our results demonstrate that 33.3% of ADS-classified C-Mi nociceptors and 40.0% of C-M nociceptors are activated by formalin during the phase 1 period in our experimental set up. The mechanism behind this activation is proposed to be through direct TRPA1 activation (Fischer et al., 2014, 2015; McNamara et al., 2007) and/or the cytotoxic effects of formalin on the nerve terminals (Hoffmann et al., 2022; Shields et al., 2010).

The mechanism leading to phase 2 nocifensive behaviours, however, is still debated. Until recently, it was believed that central sensitisation, in combination with inflammation-mediated activity in peripheral nociceptors was the primary drive of phase 2 behaviours seen in rodents (Tjølsen et al., 1992). However, recent studies indicate that direct cellular damage by high concentrations of formalin (e.g., 2%) can induce spontaneous activity in Aδ and C-fibres *in vitro* (i.e., without inflammation) (Hoffmann et al., 2022). Here we clearly show that activity dependent slowing and functionally classified C-Mi nociceptors are activated during phase 2 of the formalin response. Specifically, C-M and C-Mi nociceptors were active at equivalent proportions of 31.6% and 31.3%, respectively, during the phase 2 period after formalin exposure. These results provide insight into the underlying mechanisms of the formalin behavioural test, which remains a commonly employed assay in pain research. Furthermore, since C-fibre activity initiates central sensitisation (LaMotte et al., 1991) and likely plays a critical role in its maintenance in pathology (Koltzenburg et al., 1994), this insight into the formalin test may enable a better understanding of and treatment development for related chronic pain conditions (Woolf, 2011).

Although the electrically-evoked latency tracks of some units were lost throughout the recordings, the majority of latency tracks were maintained, demonstrating the stability of the multi-contact microneurography technique. Additionally, the ability to collect and compare the latency tracks of several units across a single experiment provides greater confidence in the identification of abnormal activity. The lack of observed abnormalities in the C-non-nociceptors suggests the effect of formalin is specific to nociceptors.

### Formalin effect on units

Given that formalin interacts with voltage-gated sodium channels, and that activity dependent slowing is primarily a function of the expression of voltage-gated channels and pumps (Namer et al., 2015; Petersson et al., 2014; Tigerholm et al., 2014), some effect on the activity dependent slowing classifications of units might be expected. Despite this, the classifications largely held, possibly reflecting that many of the direct interactions with voltage-gated sodium channels and/or TRPA1 are likely to be resolved an hour after exposure (Fischer et al., 2014). Consequently, we can be confident in the activity dependent slowing classifications, even for the units which appeared after injection.

Destruction of the sensory nerve endings by formalin would likely impact the mechanical sensitivity of affected units. Indeed, previous studies in primary nerve afferents observed an increase in threshold or loss of mechanical sensitivity when formalin was injected directly into the receptive field, but not when injecting adjacently, which is similar to our study (Hoffmann et al., 2022; McCall et al., 1996; Puig & Sorkin, 1996). Indeed, in our own preliminary studies, injection directly into the receptive field resulted in complete obliteration of the sensory afferents, also reported in other studies (Hoffmann et al., 2022). In contrast to this theory, the subsequent inflammation consequent upon local tissue damage might increase the mechanical sensitivity of surviving units, particularly of C-Mi nociceptors, which might in turn partially contribute to the mechanical allodynia and hyperalgesia seen in the formalin behavioural test (Salinas-Abarca et al., 2017). However, we did not find evidence of altered mechanical sensitivity to support this conclusion, suggesting these nocifensive evoked behaviours are primarily driven by co-occurring central sensitisation or sensitisation of A-fibre afferents.

### Differences between sexes

We saw no differences in the saphenous C-fibre populations nor the electrophysiological responses between female and male rats. Although previous studies have indicated differences in the behavioural responses of the two sexes (Pepino et al., 2023), these differences were mediated at the spinal cord level, and our results support the idea that there were no differences at the peripheral nerve level.

### Limitations and future directions

Despite our bias towards C-fibres, we found a number of units conducting in the conduction velocity range of Aδ fibres. We did not investigate the physiological properties of these Aδ fibres, however given their known activity in the formalin test (Hoffmann et al., 2022; Puig & Sorkin, 1996), further work characterising their physiological function is warranted. Additionally, it is likely that a greater proportion of Aβ fibres are present in the saphenous nerve than reported here. When this population was present in a recording, it typically appeared as a large compound action potential, and single units could not be separated. Using smaller electrical currents to minimise the size of the stimulation artifact and/or higher stimulation frequencies might make single-unit latency tracking of Aβ fibres possible. Given the recent discoveries in Aβ subclasses, including nociceptors (Bouchatta et al., 2023; Israel et al., 2025), such a study merits attention.

A further limitation of our protocol is that a low frequency of ongoing activity on the order of ≤1 Hz is unlikely to produce a substantial disruption in the latency tracks of C-fibres (Serra et al., 2012), particularly in C-M fibres which have minimal slowing at low frequency, and as seen here in the case of the C-Mi nociceptor that had double spiking where only small perturbations in latency were seen (Fig. 5b). Thus, it is possible that a higher proportion of fibres were spontaneously active, but at a frequency under the threshold that was visible using this marking technique.

Importantly, we confirm the broad and non-specific action of formalin on sensory afferents observed *in vitro*, and specifically the contribution of C-Mi nociceptors during both phases. As seen in this study, multi-contact electrodes capturing a diverse population of single units simultaneously can provide a more holistic insight into the effects of administered drugs on individual nerve subtypes. Together, these results provide clarity regarding pre-clinical nociceptive assays and their relevance to human disease.

## Acknowledgements

Analysis code and data will be shared upon reasonable request to the author.

## Author contributions

EAA, AEP and JPD planned the study. Experiments were performed and data analysed by EAA and JPD. EAA prepared the manuscript with feedback and editing by AEP and JPD. All authors have approved the final manuscript for publication.

